## Supplementary information for "A billion years arms-race between viruses, virophages and eukaryotes"

**Table S1.** Viral species included in the final multiple sequence alignments (54 taxa) with their accession numbers and source reference.

| **Species** | **ID** | **GenBank/metagenomic accessions** | **Source** |
|---|---|---|---|
| *Maverick* (*Danio rerio*) | Proteo5 | NC_007136.7: 27671708-27687993 | (Barreat and Katzourakis, 2021) |
| *Maverick* (*Xenopus tropicalis*) | Proteo18 | NC_030686.1: 6013889-6026611 | (Barreat and Katzourakis, 2021) |
| *Maverick* (*Anolis carolinensis*) | Proteo2 | NC_014778.1: 129158247-129172346 | (Barreat and Katzourakis, 2021) |
| *Maverick* (*Xiphophorus hellerii*) | Proteo19 | QPIH01000028.1: 6019497-6036410 | (Barreat and Katzourakis, 2021) |
| *Maverick* (*Oreochromis niloticus*) | Proteo15 | NW_020327416.1: 45576-61017 | (Barreat and Katzourakis, 2021) |
| *Maverick* (*Oxygymnocypris stewartii*) | Proteo11 | QVTF01001200.1: 1425766-1443135 | (Barreat and Katzourakis, 2021) |

| | | | |
|---|---|---|---|
| *Maverick* (*Plutella xylostella*) | Group2_9 | NW_011952032.1: 283307-321039 | This work |
| *Maverick* (*Ostrinia furnacalis*) | Group2_10 | NW_021132744.1: 18414-41178 | This work |
| *Maverick* (*Diachasma alloeum*) | Group2_5 | NW_021681489.1: 105940-131438 | This work |
| *Maverick* (*Diachasma alloeum*) | Group2_6 | NW_021680987.1: 30857-54497 | This work |
| *Maverick* (*Photinus pyralis*) | Group2_7 | NW_022171288.1: 44859-60568 | This work |
| *Maverick* (*Sitophilus oryzae*) | Group2_8 | NW_022147237.1: 411147-463473 | This work |
| Metagenomic NCLDV | NCLDV_Roux3157 | Trout_Epilimnion_TBL_comb48_EPIDRAFT_1006965 | (Roux et al., 2017) |
| Metagenomic NCLDV | NCLDV_Mo545 | SRX331950.48.dc.fa_4042 | (Moniruzzaman et al., 2020) |
| Metagenomic NCLDV | NCLDV_Mo338 | SRX330942.38.dc.fa_3117 | (Moniruzzaman et al., 2020) |
| Metagenomic NCLDV | NCLDV_Mo338 | SRX330942.38.dc.fa_13478 | (Moniruzzaman et al., 2020) |
| Metagenomic NCLDV | NCLDV_Roux1456 | Mendota_contig-68000307_4926_nucleotides | (Roux et al., 2017) |
| Metagenomic NCLDV | NCLDV_Sch668 | MN740268.1 | (Schulz et al., 2020) |
| *Shrimp hemocyte iridescent virus* | — | NC_055165.1 | NCBI |
| *Infectious spleen and kidney necrosis virus* | — | NC_003494.1 | NCBI |

| | | | |
|---|---|---|---|
| *Heterosigma akashiwo virus 01* | — | NC_038553.1 | NCBI |
| *Akhmeta virus* | — | NC_055230.1 | NCBI |
| *Tupanvirus soda lake* | — | KY523104.2 | NCBI |
| *Diadromus pulchellus ascovirus 4a* | — | NC_011335.1 | NCBI |
| Metagenomic PLV | PLV_BS940 | 3300009435_____Ga0115546_1002426 | (Bellas and Sommaruga, 2021) |
| Metagenomic PLV | PLV_BS718 | Ga0115028_10000066 | (Bellas and Sommaruga, 2021) |
| Metagenomic PLV | PLV_BS13 | V563K_contig_8723_len_16581_bp | (Bellas and Sommaruga, 2021) |
| Metagenomic PLV | PLV_BS395 | ERR1823950_NODE_975_length_11499_cov_287.344855 | (Bellas and Sommaruga, 2021) |
| Metagenomic PLV | PLV_BS539 | ERX2821583_NODE_576_length_19830_cov_33.560977 | (Bellas and Sommaruga, 2021) |

| | | | |
|---|---|---|---|
| Metagenomic PLV | PLV_BS262 | HanCross_NODE_1742_length_13974_cov_48.888145 | (Bellas and Sommaruga, 2021) |
| Metagenomic PLV | Plike_4 | SAF4 | (Yutin et al., 2015) |
| Metagenomic PLV | Plike_7 | INO1 | (Yutin et al., 2015) |
| Metagenomic PLV | Plike_17 | RED1 | (Yutin et al., 2015) |
| Metagenomic PLV | Plike_1 | SAF1 | (Yutin et al., 2015) |
| Metagenomic PLV | Plike_19 | YSL1 | (Yutin et al., 2015) |
| Metagenomic PLV | Plike_25 | ACE1 | (Yutin et al., 2015) |
| Metagenomic virophage | Virophage_PE125 | 3300009070_____Ga0066256_1001637 | (Paez-Espino et al., 2019) |
| Metagenomic virophage | Virophage_PE85 | 3300009781_____Ga0116178_10003357 | (Paez-Espino et al., 2019) |
| Metagenomic virophage | Virophage_PE47 | 3300012984_____Ga0164309_10000286 | (Paez-Espino et al., 2019) |
| Metagenomic virophage | Virophage_PE169 | 3300009154_____Ga0114963_10000678 | (Paez-Espino et al., 2019) |
| Metagenomic virophage | Virophage_PE184 | 3300007609_____Ga0102945_1000484 | (Paez-Espino et al., 2019) |
| Metagenomic virophage | Virophage_Roux27 | TBH_10005622 | (Roux et al., 2017) |

| | | | |
|---|---|---|---|
| *Maverick*-related virus strain Spezl | — | NC_015230.1 | NCBI |
| Yellowstone Lake virophage 7 | — | KM502591.1 | NCBI |
| Zamilon virus | — | NC_022990.1 | NCBI |
| Yellowstone Lake virophage 5 | — | NC_028269.1 | NCBI |
| Dishui Lake virophage 6 | — | MN940573.1 | NCBI |
| Dishui Lake virophage 2 | — | MN940570.1 | NCBI |
| *Bat mastadenovirus WIV17* | — | NC_034626.1 | NCBI |
| *Murine mastadenovirus A* | — | AC_000012.1 | NCBI |
| *Murine adenovirus 2* | — | NC_014899.1 | NCBI |
| *Fowl aviadenovirus 5* | — | NC_021221.1 | NCBI |
| *Frog adenovirus 1* | — | NC_002501.1 | NCBI |
| *Snake adenovirus 1* | — | NC_009989.1 | NCBI |

**Table S2.** Distribution of root positions calculated from the MCMC posterior tree sample. The best supported position of the root was on the branch leading to virophages (53.9%), followed by NCLDVs and metagenomic PLV BS539 (27.4%). Other root positions received < 6% support. The frequencies of trees with a certain position of the root were estimated by filtering different topologies in PAUP. Number of generations = 140 million.

| **Root position** | **Number of trees** | **Percentage** | **Cumulative sum** |
|---|---|---|---|
| **Virophages** | **15099** | **53.923** | **53.923** |
| NCLDVs + PLV_BS539 | 7672 | 27.399 | 81.322 |
| PLV_BS539 | 1572 | 5.614 | 86.936 |
| NCLDVs | 1489 | 5.318 | 92.254 |
| Virophages + NCLDVs + PLV_BS539 | 1388 | 4.957 | 97.211 |
| PLV_BS13 | 185 | 0.661 | 97.872 |
| Virophages + PLV_BS13 | 159 | 0.568 | 98.439 |
| Adenoviruses | 107 | 0.382 | 98.821 |
| Virophages + NCLDVs | 44 | 0.157 | 98.979 |
| Virophages + NCLDVs + PLV_BS539 + PLV_BS13 | 35 | 0.125 | 99.104 |
| Adenovirus + NCLDVs + PLV_BS539 | 29 | 0.104 | 99.207 |
| Adenoviruses + NCLDVs | 4 | 0.014 | 99.221 |
| Virophages + Adenoviruses | 3 | 0.011 | 99.232 |
| NCLDVs + PLV_BS539 + most PLVs | 1 | 0.004 | 99.236 |

| | | | |
|---|---|---|---|
| Virophages + PLV_BS13 + Adenoviruses | 1 | 0.004 | 99.239 |
| Adenovirus + PLV_BS13 | 1 | 0.004 | 99.243 |
| most PLVs | 0 | 0.000 | 99.243 |
| *Mavericks 1* + *Mavericks 2* | 0 | 0.000 | 99.243 |
| *Mavericks 1* | 0 | 0.000 | 99.243 |
| *Mavericks 2* | 0 | 0.000 | 99.243 |
| Virophages + most PLVs | 0 | 0.000 | 99.243 |
| Adenoviruses + most PLVs | 0 | 0.000 | 99.243 |
| *Mavericks 1* + *Mavericks 2* + PLV_BS13 | 0 | 0.000 | 99.243 |
| *Mavericks 1* + *Mavericks 2* + Adenoviruses + PLV_BS13 | 0 | 0.000 | 99.243 |
| Others | 212 | 0.757 | 100 |

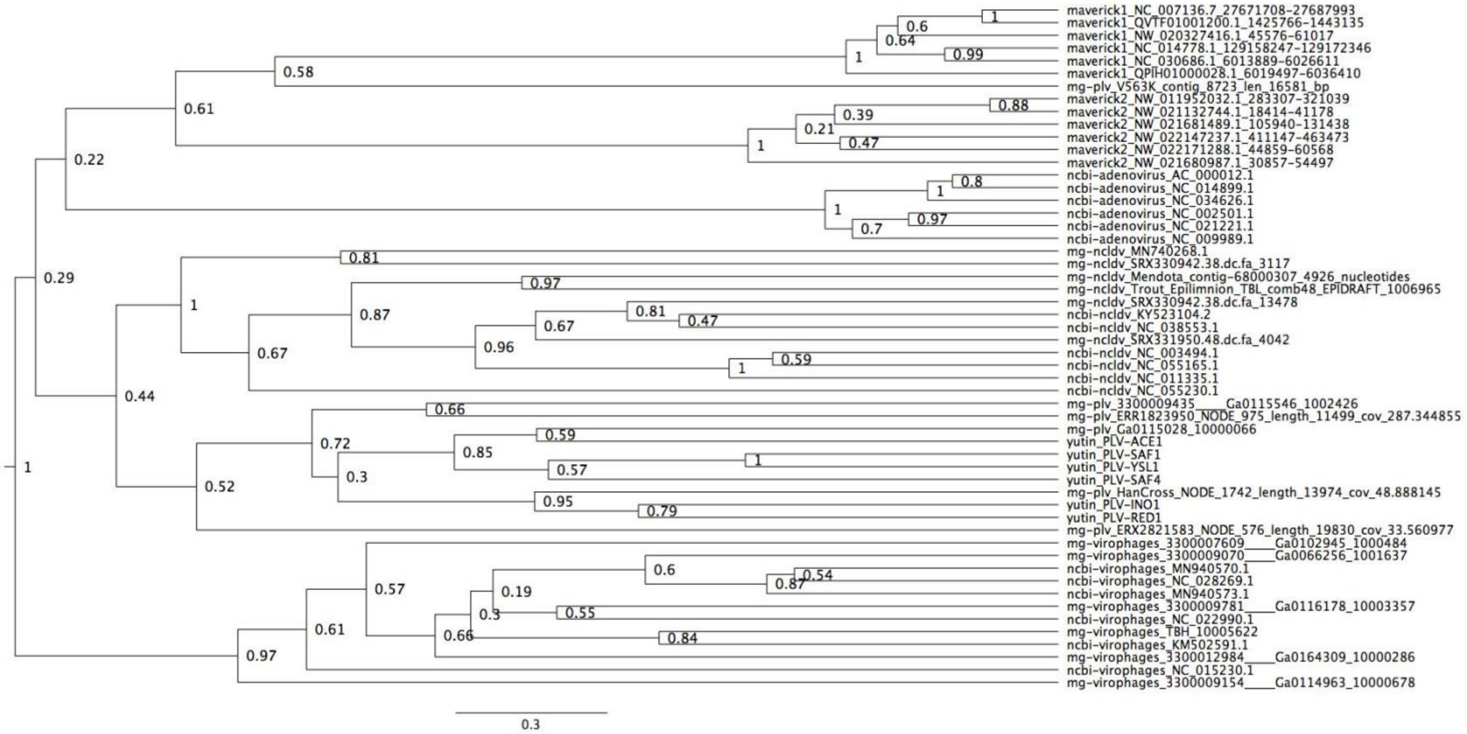


**Figure S1.** Bayesian maximum credibility tree of the major capsid protein inferred with a relaxed molecular clock. Virophages are monophyletic with a high posterior support (0.97). The root falls between virophages and all other elements. Tree estimated from 200 million MCMC generations and a 25% relative burn-in.

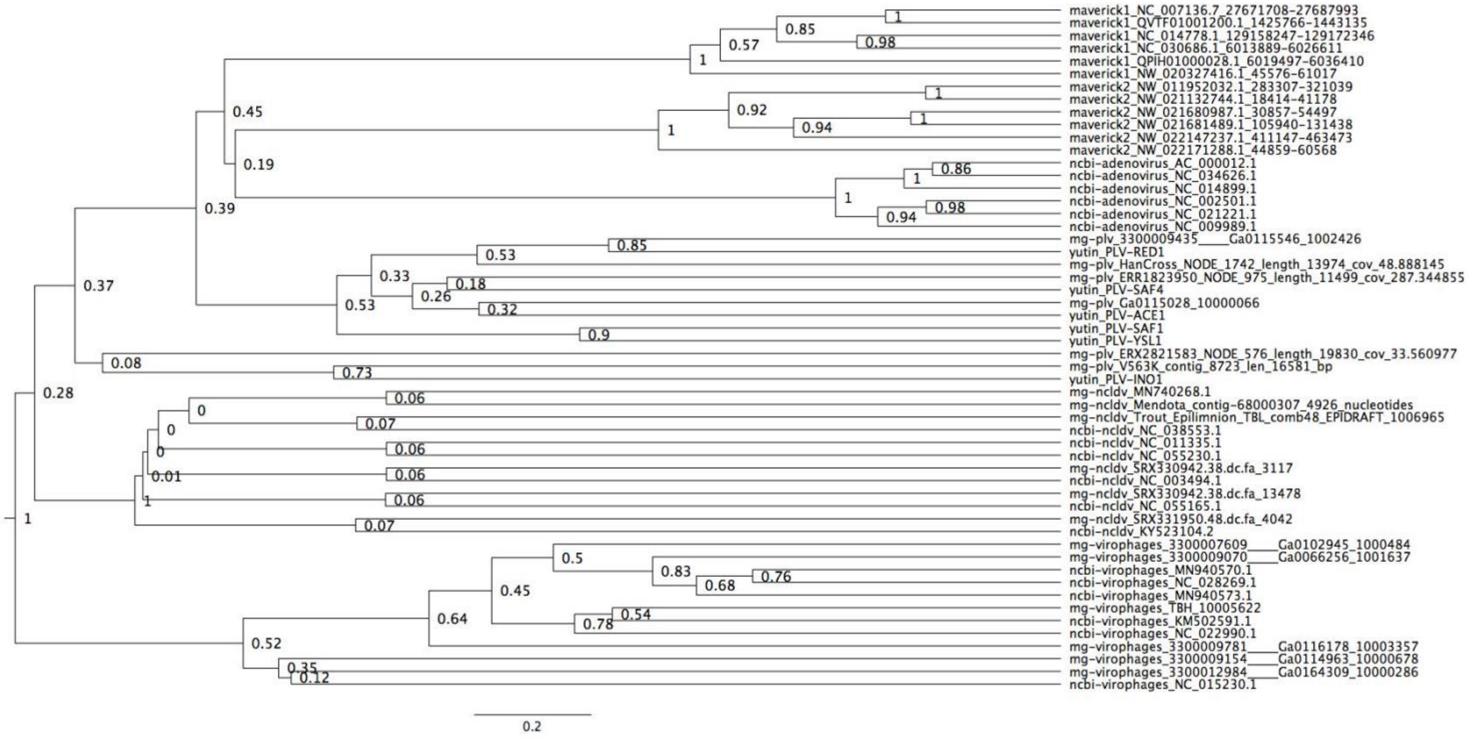


**Figure S2.** Bayesian maximum credibility tree of the minor capsid protein inferred with a relaxed molecular clock. Virophages are monophyletic with a moderate posterior support (0.52). The root falls between virophages and all other elements. Tree estimated from 200 million MCMC generations and a 25% relative burn-in.

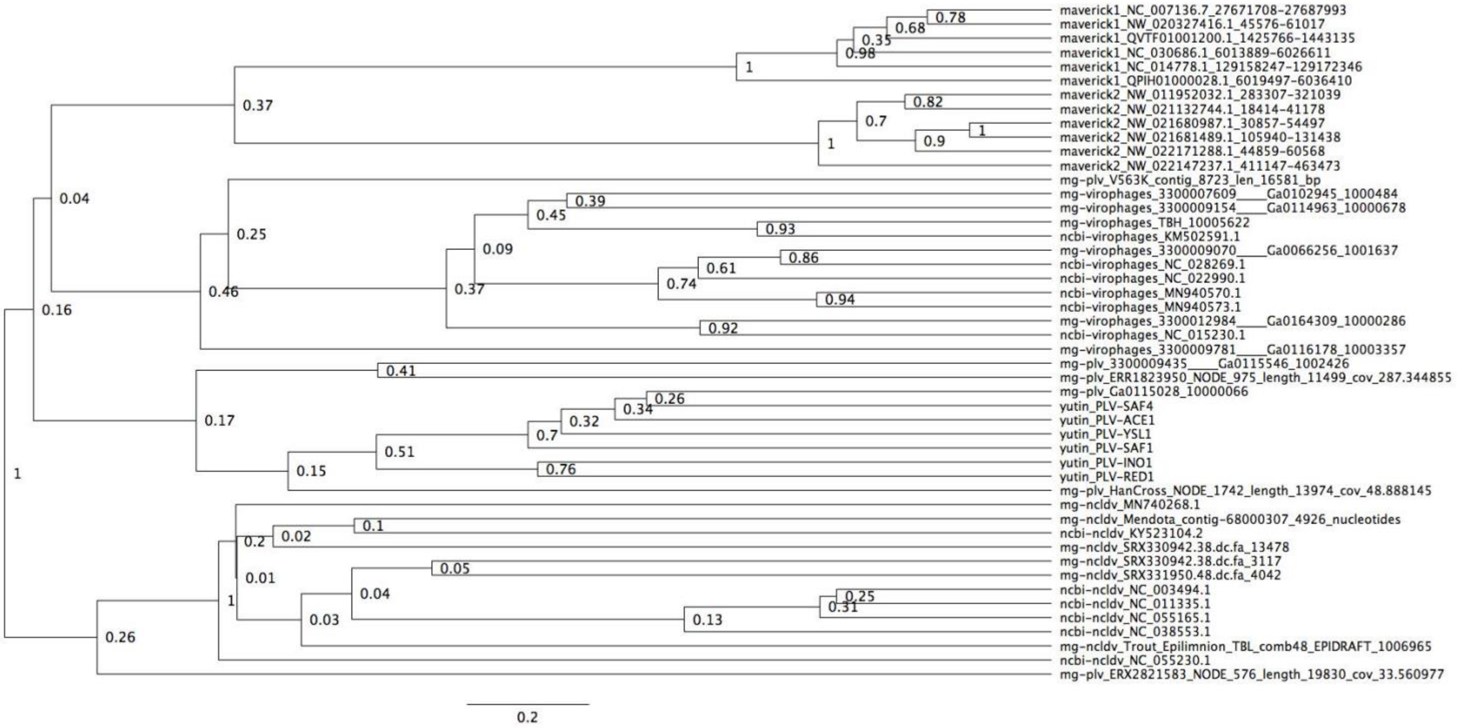


**Figure S3.** Bayesian maximum credibility tree of the ATPase inferred with a relaxed molecular clock. We did not use a monophyletic constraint on virophages in this tree. The root falls between a clade formed by NCLDVs plus a metagenomic PLV (0.26 posterior probability), and all other elements. According to this tree, virophages are not monophyletic. Tree estimated from 200 million MCMC generations and a 25% relative burn-in. Adenoviruses are not included in this tree since they encode a non-homologous ATPase of the ABC superfamily (Burroughs, et al., 2007).

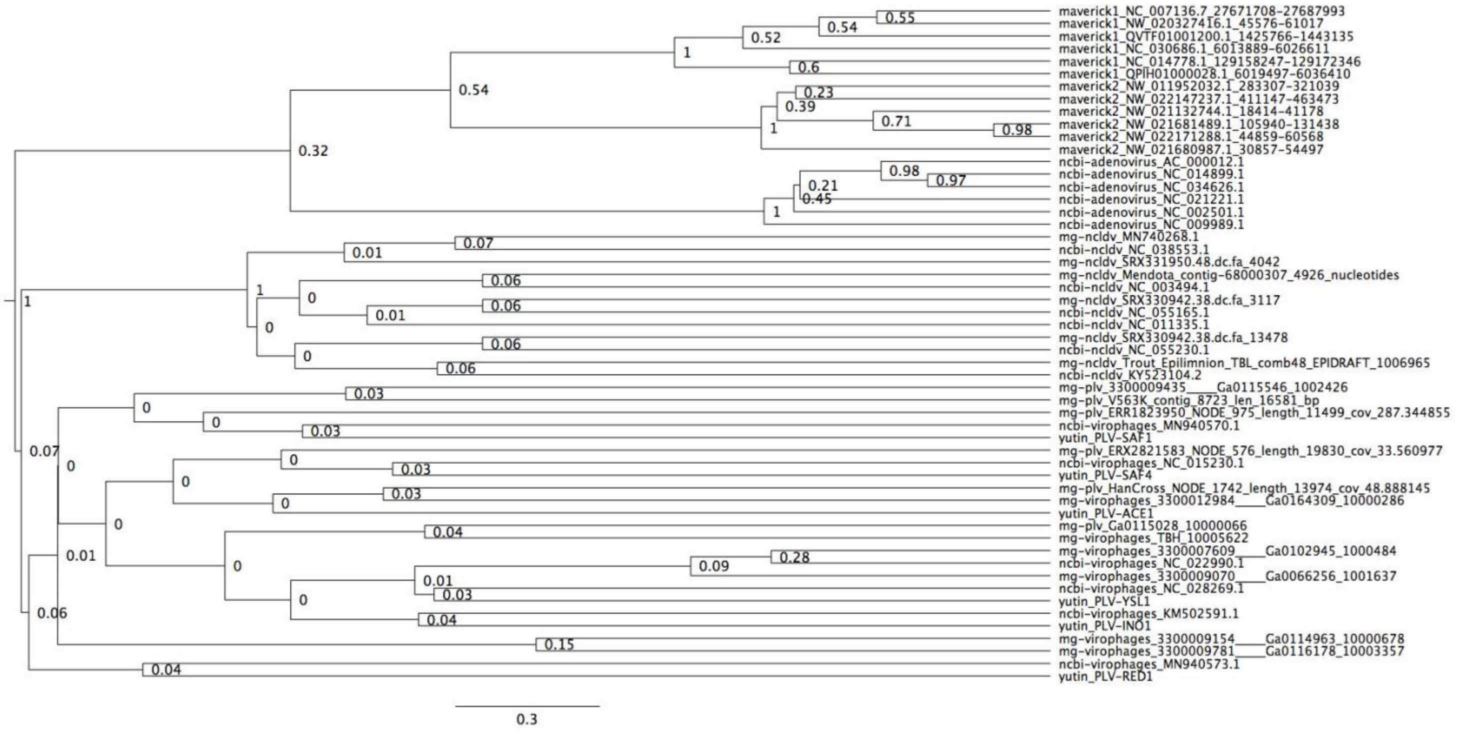


**Figure S4.** Bayesian maximum credibility tree of the protease inferred with a relaxed molecular clock. We did not use a monophyletic constraint on virophages in this tree. The root falls between a clade formed by adenoviruses and *Mavericks* and all other elements. However, notice how virophages do not form a monophyletic group. Tree estimated from 200 million MCMC generations and a 25% relative burn-in.

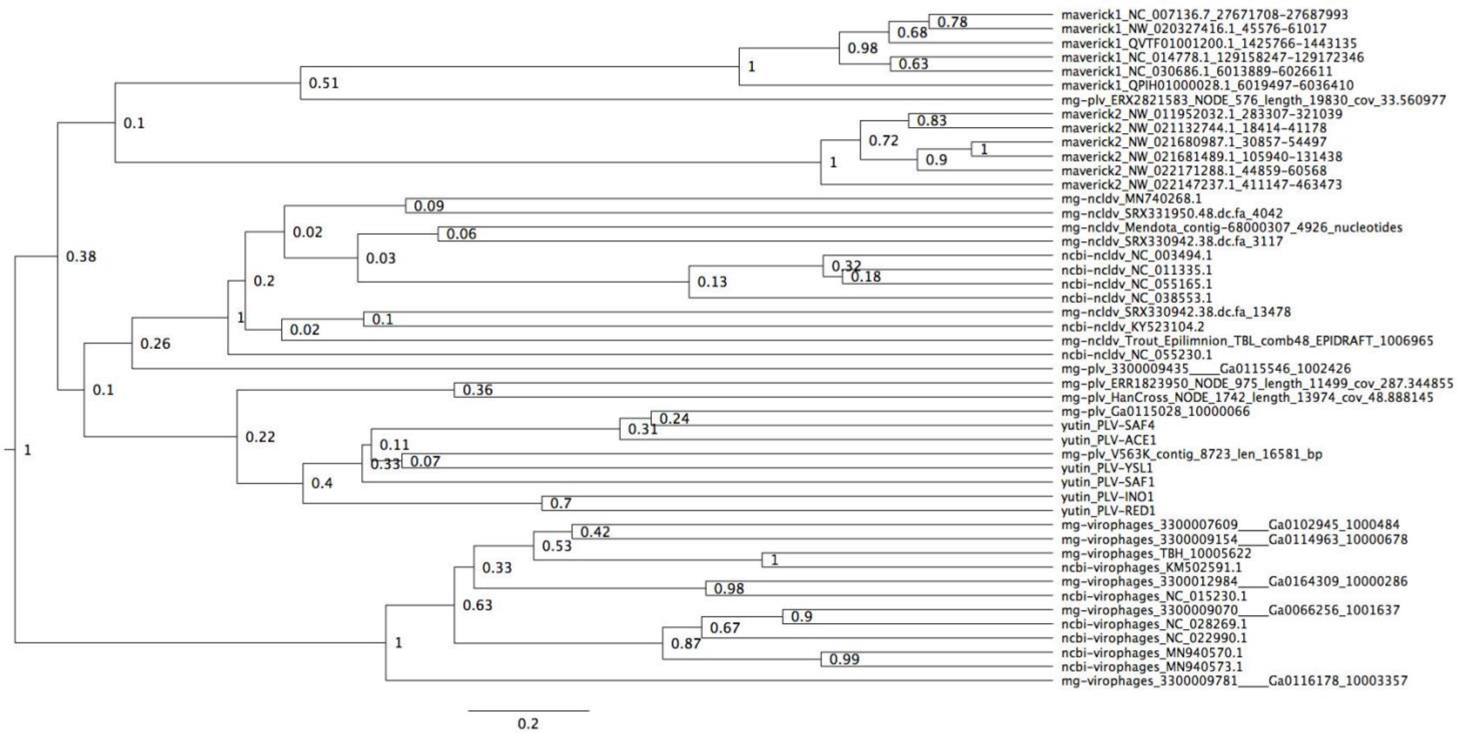


**Figure S5.** Bayesian maximum credibility tree of the ATPase inferred with a relaxed molecular clock, and using a monophyletic constraint on virophages. The root of this tree falls between virophages and all other elements. Tree estimated from 200 million MCMC generations and a 25% relative burn-in. Adenoviruses are not included in this tree since they encode a non-homologous ATPase of the ABC superfamily (Burroughs, et al., 2007).

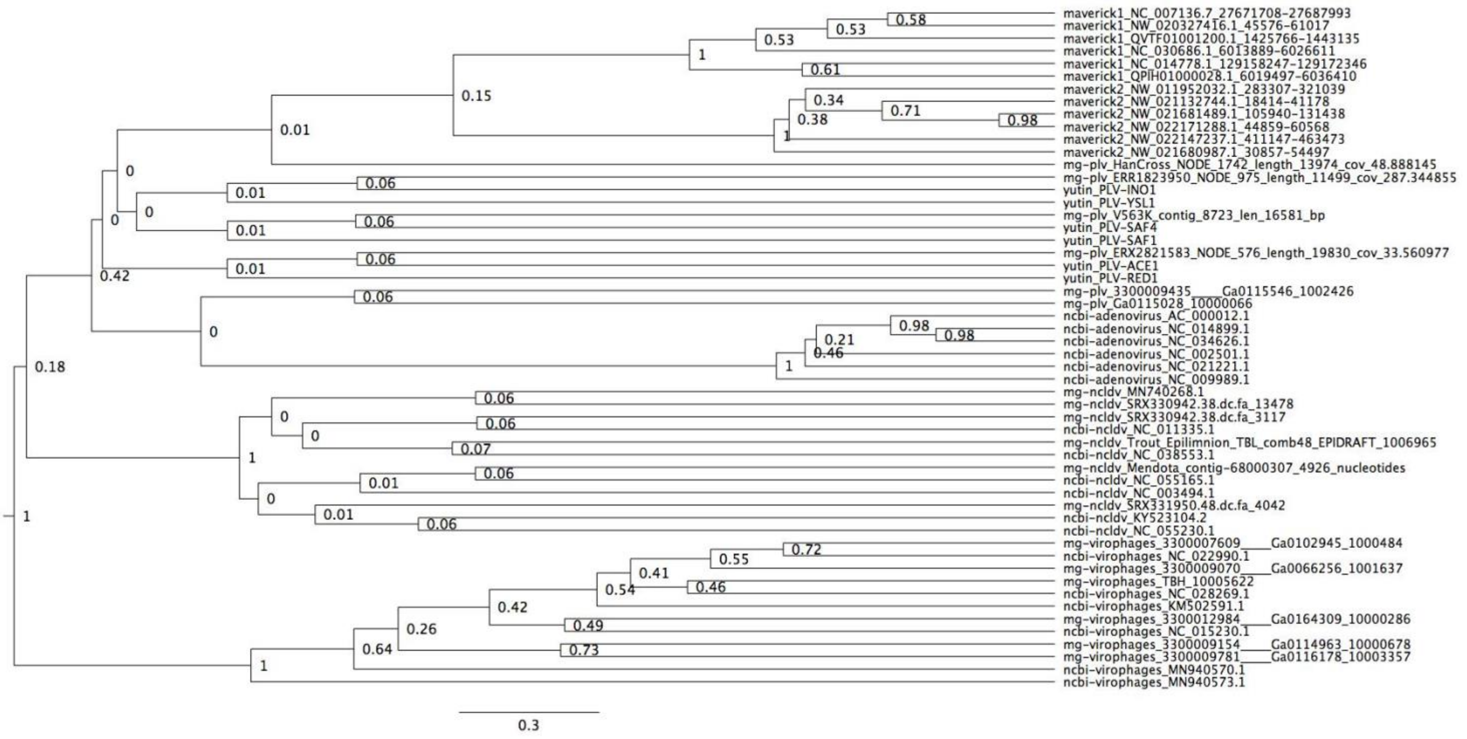


**Figure S6.** Bayesian maximum credibility tree of the protease inferred with a relaxed molecular clock, and using a monophyletic constraint on virophages. The root falls between virophages and all other elements. Tree estimated from 200 million MCMC generations and a 25% relative burn-in.

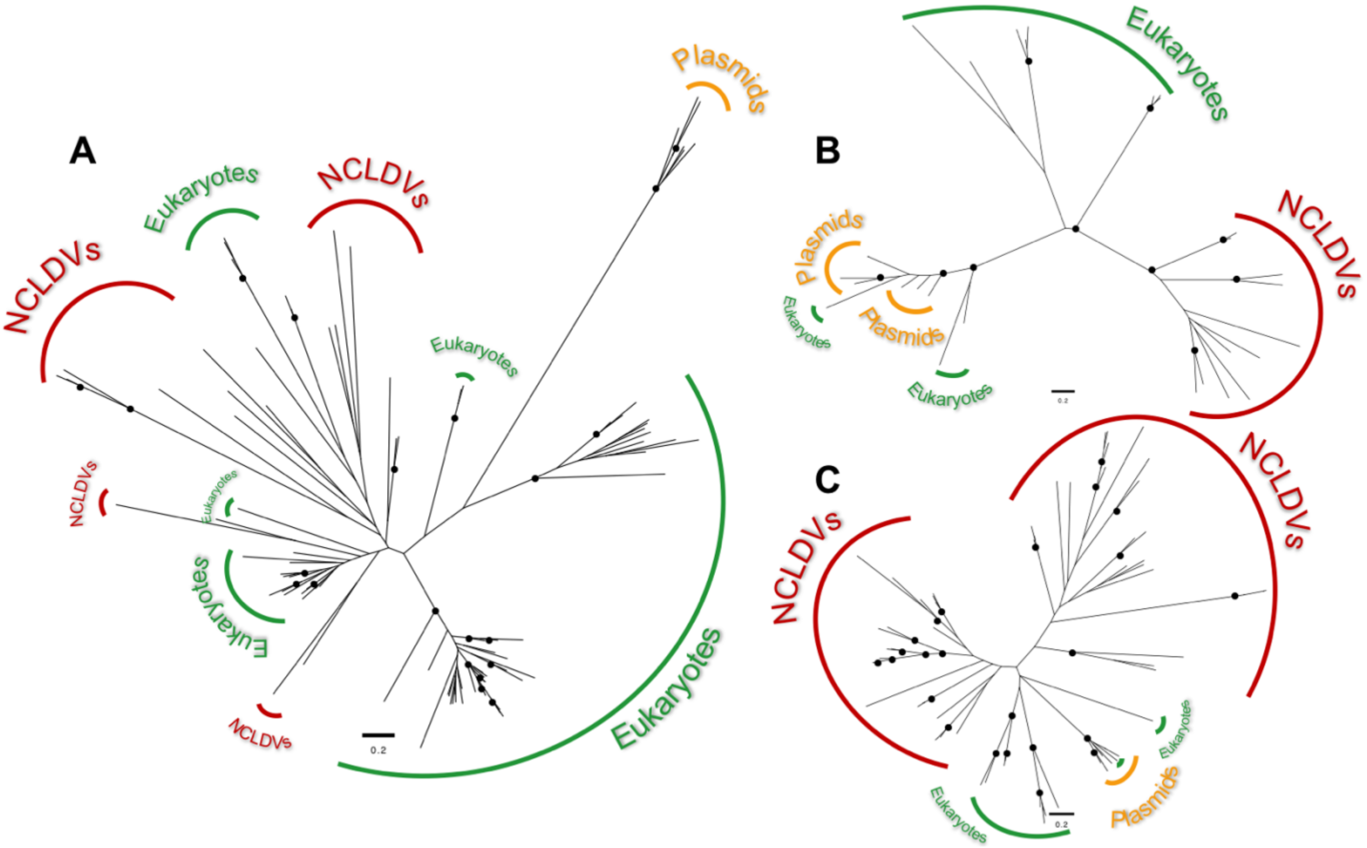


**Figure S7.** Maximum-likelihood unrooted phylogenetic trees of the transcriptional homologues encoded in cytoplasmic linear plasmids. (A) Trees for the DNA-directed RNA polymerase II, subunit Rpb2, (B) mRNA capping enzyme, and (C) helicase. In all cases, the topologies rule out a sister grouping of the NCLDV and cytoplasmic linear plasmid homologues, suggesting they were acquired independently. Black circles indicate bootstrap support ≥ 0.94 after 1,000 replicates.

**Table S3.** Cytoplasmic linear plasmids used for querying the databases in search for protein homologues.

| **Plasmid** | **GenBank accession** | **Host organism** (Order Saccharomycetales) |
|---|---|---|
| pDH4C | MF795093.1 | *Debaryomyces hansenii* (Debaryomycetaceae) |
| pGK12 | X07776.1 | *Kluyveromyces lactis* (Saccharomycetaceae) |
| pKPGS115 | CP014724.1 | *Komagataella phaffii* (Phaffomycetaceae) |
| pKP | CP014714.1 | *Komagataella phaffii* (Phaffomycetaceae) |
| pKPCBS743 | MG491503.1 | *Komagataella phaffii* (Phaffomycetaceae) |
| pPac1-1 | AM180622.1 | *Millerozyma acacia* (Debaryomycetaceae) |
| pPE1B | AJ278986.2 | *Schwanniomyces etchellsi* (Debaryomycetaceae) |
| pSKL | X54850.1 | *Lachancea kluyveri* (Saccharomycetaceae) |
